# Distinct visual processing networks for foveal and peripheral visual fields

**DOI:** 10.1101/2024.06.24.600415

**Authors:** Jie Zhang, Huihui Zhou, Shuo Wang

## Abstract

Foveal and peripheral vision are two distinct modes of visual processing essential for navigating the world. However, it remains unclear if they engage different neural mechanisms and circuits within the visual attentional system. Here, we trained macaques to perform a free-gaze visual search task using natural face and object stimuli and recorded a large number of 14588 visually responsive neurons from a broadly distributed network of brain regions involved in visual attentional processing. Foveal and peripheral units had substantially different proportions across brain regions and exhibited systematic differences in encoding visual information and visual attention. The spike-LFP coherence of foveal units was more extensively modulated by both attention and visual selectivity, thus indicating differential engagement of the attention and visual coding network compared to peripheral units. Furthermore, we delineated the interaction and coordination between foveal and peripheral processing for spatial attention and saccade selection. Finally, the search became more efficient with increasing target-induced desynchronization, and foveal and peripheral units exhibited different correlations between neural responses and search behavior. Together, the systematic differences between foveal and peripheral processing provide valuable insights into how the brain processes and integrates visual information from different regions of the visual field.

**Significance Statement:** This study investigates the systematic differences between foveal and peripheral vision, two crucial components of visual processing essential for navigating our surroundings. By simultaneously recording from a large number of neurons in the visual attentional neural network, we revealed substantial variations in the proportion and functional characteristics of foveal and peripheral units across different brain regions. We uncovered differential modulation of functional connectivity by attention and visual selectivity, elucidated the intricate interplay between foveal and peripheral processing in spatial attention and saccade selection, and linked neural responses to search behavior. Overall, our study contributes to a deeper understanding of how the brain processes and integrates visual information for active visual behaviors.

## Introduction

Visual attention acts as the gateway to cognitive processing by enabling the brain to selectively prioritize specific stimuli for further processing [1–3]. A significant body of literature has extensively documented the neural networks, pathways, and dynamics that govern the brain’s selection, processing, and integration of information to accomplish specific objectives [4, 5]. Visual attention is crucial for active visual search [6], and previous studies have elucidated its neural mechanisms, involving a complex interplay of various brain regions, including area V4 [7–10], inferotemporal (IT) cortex [11–13], and lateral prefrontal cortex (LPFC) [14]. These regions coordinate attentional resources and establish intricate networks that dynamically adjust sensory processing in response to the goals and intentions of the animal [15]. However, previous studies have predominantly focused on the peripheral receptive fields (RFs), with very few investigating foveal processing of attention in these brain areas [16, 17]. It remains largely unknown whether foveal and peripheral units have different response profiles and are differentially engaged in the attention network. Investigating foveal processing will provide critical new information for a comprehensive understanding of visual attentional processing in these brain areas.

Due to the limited capacity of attention [18–20], the relationship between central and peripheral visual fields in the attentional process is intricate. Paying attention to peripheral regions reduces the visual processing of foveated objects [21], and an increase in demand for the central task decreases the response of brain regions involved in processing task-irrelevant stimuli in the background [22]. Behaviorally, irrelevant peripheral stimuli influence the performance of tasks presented in the central region [23, 24], and better performance on tasks in the central region is associated with enhanced suppression of distracting irrelevant information in the periphery [25]. Concurrent attention to competing features in the central and peripheral visual fields belonging to the same visual modality results in significant interference, which is not the case when features belong to different visual modalities [26]. Both central [27–30] and peripheral [28, 31] visual fields play a crucial role in visual search, masking which results in longer search times and lower accuracy. However, it remains unclear whether and how foveal and peripheral neurons in attention-processing brain areas coordinate with each other during search behaviors.

In addition to modulating response gain [4, 32, 33], attention influences the coherence between spike and local field potential (LFP) within V4 [8, 13, 34] and between V4 and IT [13]. Primates actively explore their environment through saccadic eye movements, often accompanied by brain activity in the theta frequency band [35]. In particular, both classic studies [34] and our own prior studies [36] have shown desynchronization for attended stimuli in the theta frequency band. Therefore, in this study, we hypothesize that different brain areas involved in visual attention coordinate and interact with each other through spike-LFP coherence in the theta frequency band. Furthermore, we hypothesize that neurons with a foveal RF exhibit different theta coherence compared to neurons with a peripheral RF. In particular, we examined whether theta desynchronization [34, 36] within and between brain areas predicted search behaviors.

In conjunction with visual attention, visual category selectivity is a hallmark of the IT cortex [37–39]. Additionally, visual category selectivity, especially face selectivity [40], is notably evident in the orbitofrontal cortex (OFC). While classic studies show that the OFC plays a critical role in value-based decision making [41], it has also been implicated in attentional processing [42–44]. For example, studies using value-related visual stimuli during free viewing have shown that the farther the gaze point is from the value stimulus, the smaller the neuronal response in the OFC, indicating that overt attention regulates OFC activity [43]. OFC neuronal activity reflects the value of newly appeared stimuli that attract attention while reducing the representation of old stimuli, showing that covert attention affects stimulus value representation [42]. Moreover, bottom-up attention modulates OFC activity to reflect the value of the attended stimulus [44]. In particular, the OFC receives visual input from the IT cortex, and the face-selective regions of the IT cortex receive projections from the OFC [45–47]. Such reciprocal connections prompt questions about how prefrontal areas like the OFC interact with classical brain areas of visual attention in the temporal cortex. In this study, we hypothesize that the OFC is also a critical part of the feature attention network. To test this hypothesis, we investigated the role of the OFC in both feature attention and category selectivity, in the context of foveal and peripheral processing at both the neuronal and neural circuit levels.

Together, this study aims to address the overarching hypothesis that foveal units have different response profiles and functional connectivity compared to peripheral units. Specifically, we hypothesize that (1) different brain areas consist of different proportions of foveal and peripheral units, (2) the neural firing rate for category selectivity and feature attention differs between foveal and peripheral units, (3) spike-LFP coherence in the theta frequency band differs between foveal and peripheral units when encoding visual categories and attention, and (4) theta desynchronization predicts search behaviors. To test these hypotheses, we simultaneously recorded units with foveal or peripheral RFs in V4, IT, LPFC, and OFC while monkeys performed a category-based free-gaze visual search task. Our study represents one of the first to investigate foveal processing in these brain areas and offers novel insights into neural processing in the central and peripheral visual fields during active vision.

## Results

### Behavior

Two monkeys performed a free-gaze visual search task (**Fig. 1A**; see **Methods**), where their objective was to fixate on one of the two search targets that matched the same category as the cue. Both monkeys performed the task proficiently, with accuracy rates of 91.78%±1.83% (mean±SD across sessions) for monkey S and 85.23%±3.66% for monkey E. Monkeys had a similar accuracy in finding faces (89.19% ±4.11%) versus houses (89.31%±4.29%). The mean reaction time (RT), from the onset of the search array to the onset of the last fixation, was 411.47±67.01 ms (mean±SD across sessions; face-target trials: 398.66±76.75 ms, house-target trials: 423.94±60.73 ms; two-tailed paired *t*-test across sessions: *t*(174) = 10.10, P = 3.55×10^−19^). The mean fixation duration was 208.24±153.77 ms (mean±SD across fixations). We found that fixations on targets was longer than fixations on distractors (two-way ANOVA [target versus distractor × face versus house]; main effect of target versus distractor: *F*(1,79622) = 9908.87, P < 10^−50^), and fixations on faces was longer than fixations on houses (main effect of face versus house: *F*(1,79622) = 43.78, P = 3.70×10^−11^; interaction: *F*(1,79622) = 47.51, P = 5.52×10^−12^). Consistent with a longer RT for houses, there were more fixations when monkeys searched for houses compared to faces (P = 1.75×10^−135^).

**Fig. 1.**
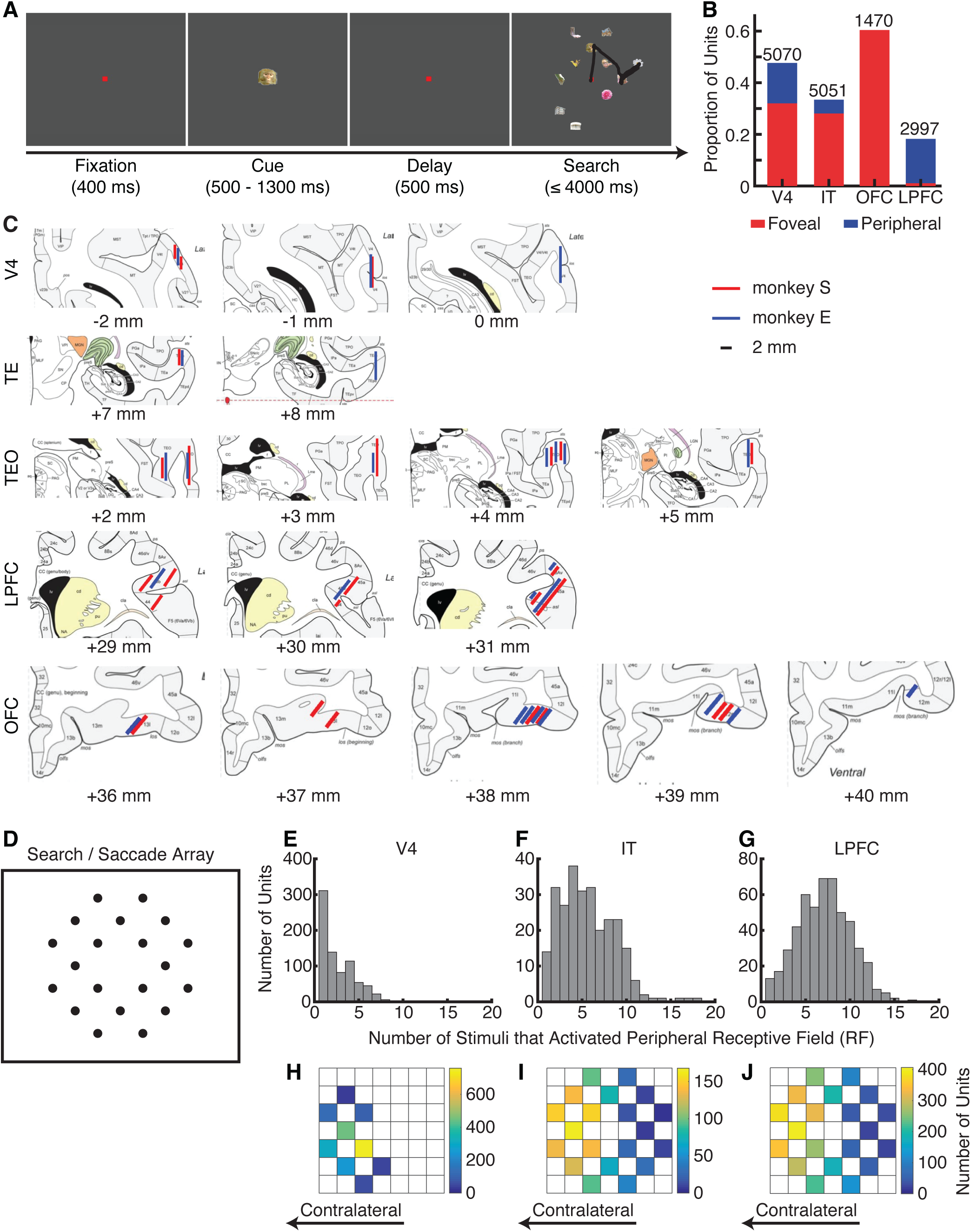
Task and summary of units. **(A)** Task. Monkeys initiated the trial by fixating on a central point for 400 ms. A cue was then presented for 500 to 1300 ms. After a delay of 500 ms, the search array with 11 items appeared. Monkeys were required to fixate on one of the two search targets that belonged to the same category as the cue for at least 800 ms to receive a juice reward. **(B)** The proportion of foveal and peripheral units in each brain area. Red: units with a focal foveal receptive field. Blue: units with a peripheral receptive field. The total number of visually responsive units is indicated at the top of each bar. **(C)** Recording sites in two monkeys overlaid on the atlas of the rhesus monkey brain in stereotaxic coordinates [69]. The red and blue lines represent the estimated spatial range of recordings in monkey S and monkey E, respectively. The numbers below indicate the rostral (+) or caudal (-) distances of the slices from the infraorbital ridge (Ear Bar Zero). **(D)** The 20 possible stimulus locations in the visual search task and the visually guided saccade task. Peripheral receptive fields (RFs) were mapped using the visually guided saccade task, which had the same 20 possible stimulus locations as the visual search task. **(E-G)** Histogram of the number of stimuli that activated the peripheral RF. **(H-J)** The aggregated tuning regions of the peripheral units with a localized RF. Color bars show the number of units with tuning regions in a given stimulus location. The right side of the brain was recorded for both monkeys. **(E, H)** V4 units. **(F, I)** IT units. **(G, J)** LPFC units.

Importantly, we found that the search accuracy (**Fig. S1A**; one-way ANOVA; P = 0.25), RT (**Fig. S1B**; P = 0.89), and fixation duration (**Fig. S1C**; P = 0.70) did not differ significantly between sessions with recordings from different brain regions. Therefore, the differences across brain regions could not be attributed to behavioral differences across sessions.

### Units with foveal versus peripheral receptive fields

We recorded a total number of 6871 units from area V4, 8641 units from the IT cortex (including both TE and TEO), 5622 units from the OFC, and 9916 units from the LPFC (see **Fig. 1C** for detailed recording locations). 5070 units from area V4, 5051 units from the IT cortex, 1470 units from the OFC, and 2997 units from the LPFC had a significant visually evoked response (i.e., the response to the cue or search array was significantly greater than the response to the baseline; Wilcoxon rank-sum test: P < 0.05). Among these visually responsive units, 1624 units from V4, 1419 units from the IT cortex, 888 units from the OFC, and 32 units from the LPFC had a focal foveal RF (see **Methods**; **Fig. 1B**), while 781 units from V4, 268 units from the IT cortex, no units from the OFC, and 514 units from the LPFC had a localized peripheral RF (**Fig. 1B**; the rest either had a broad foveal RF or an unlocalized peripheral RF; see **Methods**). Therefore, the proportion of foveal versus peripheral units varied substantially across brain areas. In this study, we specifically compared units with a focal foveal RF and units with a peripheral RF.

We first characterized the peripheral RFs. We found that the size of the peripheral RF (quantified by the number of stimuli that activated the unit; **Fig. 1D**) increased from V4 (2.72±2.03 [mean±SD] stimuli; **Fig. 1E, H**) to IT (5.62±3.07 stimuli; **Fig. 1F, I**) to LPFC (7.11±2.95 stimuli; **Fig. 1G, J**; two-tailed two-sample *t*-test across units: V4 versus IT: *t*(1047) = 17.51, P = 1.95×10^−60^; V4 versus LPFC: *t*(1293) = 31.75, P = 5.03×10^−164^; IT versus LPFC: *t*(780) = 6.63, P = 6.35×10^−11^). Qualitatively, we found that some parts of the V4 RFs were more densely represented by our recorded neuronal population (**Fig. 1H**), while IT and LPFC RFs were more homogeneously represented along the horizontal positions (**Fig. 1I, J**). Additionally, in the IT cortex and LPFC, the vertical positions were less represented than the horizontal positions, but we did not observe a decay as a function of distance from the fovea (**Fig. 1I, J**).

### Differences in feature attention effect and category selectivity between foveal and peripheral units

We analyzed the feature attention effect and category selectivity in each brain area, separately for foveal and peripheral units (note that the OFC had no peripheral units whereas the LPFC only 32 foveal units). It is worth noting that we matched the exact stimulus between targets and distractors for the firing rate analysis. For both the firing rate and coherence analyses of peripheral units, we ensured that there was no impending saccade toward the stimulus in the peripheral RF, consistent with prior studies on feature-based attention [12].

We identified *category-selective units* that differentiated fixations on faces and houses and *target-selective units* that differentiated fixations on targets and distractors (**Methods**). For foveal units, we found that V4 (35.16%; binomial P < 10^−20^; **Fig. 2A, D**), IT (60.61%; binomial P < 10^−20^; **Fig. 2B, E**), and OFC (74.21%; binomial P < 10^−20^; **Fig. 2C, F**) all had an above-chance population of category-selective units, suggesting that foveal units in these brain areas encoded visual category information (see **Fig. S2A-C** for alignment at the cue onset and **Fig. S2G-I** for response in the visually guided saccade task). Similarly, V4 (21.43%; binomial P < 10^−20^; **Fig. 2A, H**), IT (25.02%; binomial P < 10^−20^; **Fig. 2B, I**), and OFC (17.23%; binomial P < 10^−20^; **Fig. 2C, J**) all had an above-chance population of target-selective units, suggesting that foveal units in these brain areas also encoded visual attention. Interestingly, category-selective units were not more likely to be target-selective units in V4 (χ^2^ test between the proportion of target-selective units within category-selective units [*n*_Both_ / *n_Category-Selective_*] versus the proportion of target-selective units within all units [*n*_Target-Selective_ / *n*_All_]: P = 0.994), IT (P = 0.75), or OFC (P = 0.78), suggesting that category-selective and target-selective units were largely distinct populations. In addition, when encoding visual category information, foveal units in both V4 and IT had a significantly faster response latency than those in the OFC (see **Methods**; **Fig. 2G**; V4: 79.91±45.89 ms [mean±SD across units]; IT: 76.48±40.94 ms; OFC: 92.48±53.12 ms; two-tailed two-sample *t*-test: V4 versus OFC: *t*(1280) = 4.39, P = 1.21×10^−5^; IT versus OFC: *t*(1335) = 6.05, P = 1.86×10^−9^; V4 versus IT: *t*(1751) = 1.65, P = 0.098; Bonferroni correction for multiple comparisons). When encoding visual attention, foveal units in the OFC had a significantly faster response latency than those in the IT cortex (**Fig. 2K**; V4: 70.55±59.46 ms; IT: 76.94±58.73 ms; OFC: 63.08±54.85 ms; IT versus OFC: *t*(1036) = 3.75, P = 1.87×10^−4^; V4 versus IT: *t*(1389) = 2.01, P = 0.04; V4 versus OFC: *t*(1109) = 2.04, P = 0.04).

**Fig. 2.**
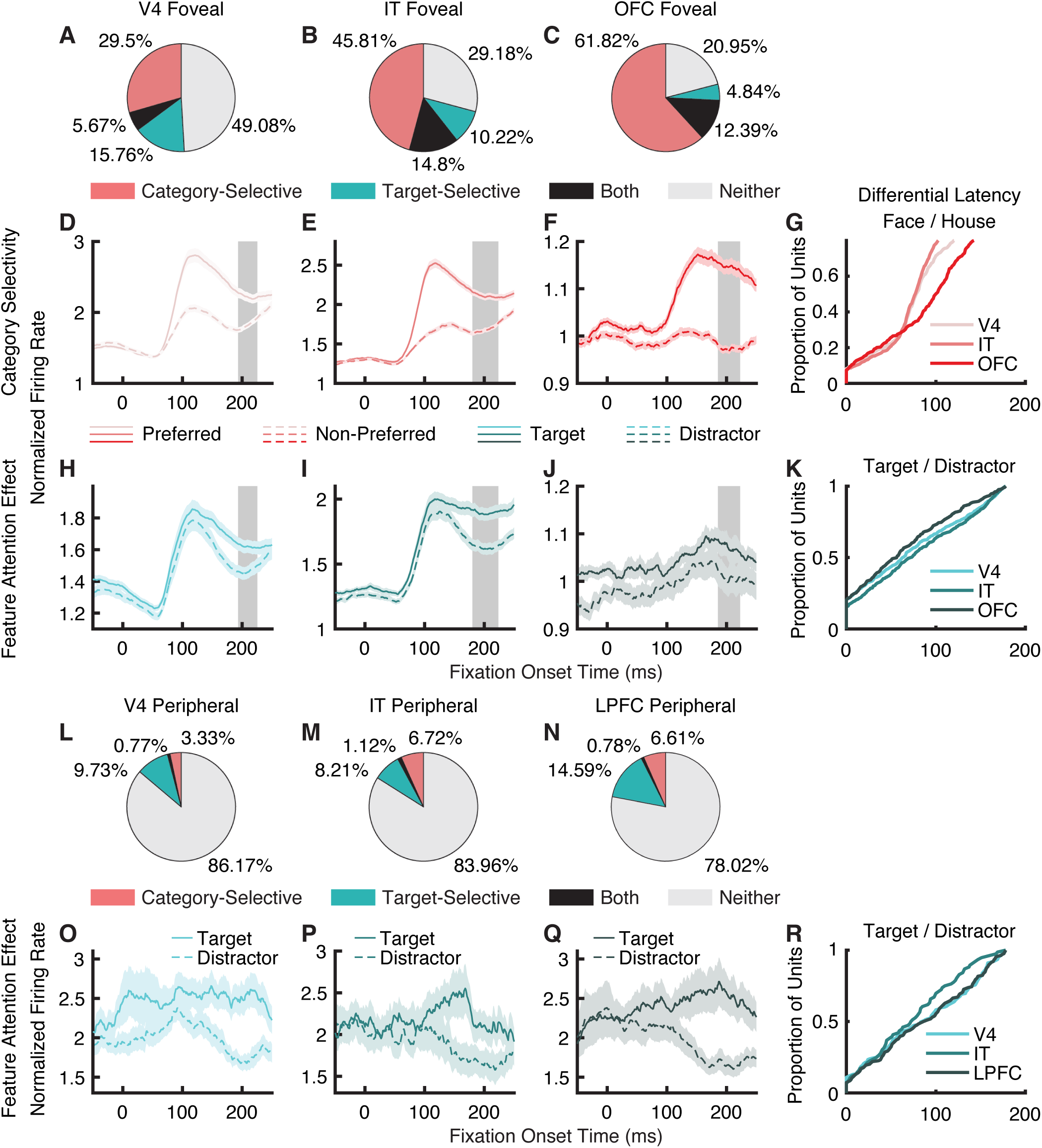
Feature attention effect and category selectivity in units with focal foveal versus peripheral receptive fields (RFs). **(A-K)** Units with a focal foveal RF. **(L-R)** Units with a peripheral RF. **(A-C, L-N)** Population summary of target-selective and category-selective units. **(D-G)** Category selectivity. **(H-K, O-R)** Feature attention effect. **(D-F, H-J, O-Q)** Shown are normalized firing rates. Shaded area around each line denotes ±SEM across units. The gray shade indicates the average time of fixation before the next saccade (mean±SD across sessions). **(G, K, R)** Differential latency. The cumulative distribution of differential latencies computed individually for each brain area. **(A, D, H, L, O)** V4. **(B, E, I, M, P)** IT. **(C, F, J, N, Q)** TEO.

For peripheral units, V4 (4.10%; binomial P = 0.86; **Fig. 2L**), IT (7.84%; binomial P = 0.016; **Fig. 2M**), and LPFC (7.39%; binomial P = 0.0072; **Fig. 2N**) only had a small population of category-selective units, suggesting that peripheral units in these brain areas barely encoded visual category information. Although V4 (10.5%; binomial P = 1.75×10^−10^; **Fig. 2L, O**), IT (9.33%; binomial P = 0.001; **Fig. 2M, P**), and LPFC (15.37%; binomial P < 10^−20^; **Fig. 2N, Q**) had an above-chance population of target-selective units, the percentage was significantly lower compared to foveal units (χ^2^ test; V4: P = 5.75×10^−11^; IT: P = 1.71×10^−8^; see **Fig. S2D-F** for alignment at the array onset). In contrast to foveal units (**Fig. 2K**), peripheral units in these brain areas did not show a significant difference in response latency when encoding visual attention (**Fig. 2R**; V4: 85.10±58.35 ms; IT: 73.75±49.56 ms; LPFC: 86.29±56.33 ms; V4 versus IT: *t*(275) = 1.50, P = 0.13; V4 versus LPFC: *t*(363) = 0.20, P = 0.84; IT versus LPFC: *t*(238) = 1.67, P = 0.097). Notably, foveal units responded significantly faster than peripheral units for feature attention effect in V4 (two-tailed two-sample *t*-test: *t*(617) = 2.26, P = 0.02; but not in IT: *t*(512) = 0.61, P = 0.54).

Together, foveal and peripheral units exhibited systematic differences in encoding visual information and visual attention.

### Different functional connectivity for foveal versus peripheral units

Are foveal and peripheral units engaged in the same functional network? To answer this question, we analyzed the coherence between spikes and LFPs recorded simultaneously across brain areas (see **Methods**). For each type of units, we included spikes from brain areas with at least 20 units and LFPs from all four brain areas. Consistent with previous studies [34, 36], spike-LFP coherence was enhanced in the theta frequency band. Therefore, we focused on theta coherence in this study.

We first investigated the modulation by attention by examining the change in synchronization in the attended (fixations on targets) stimuli relative to the non-attended stimuli (fixations on distractors). We found that for both foveal and peripheral units (regardless of target selectivity based on firing rate), spikes desynchronized with LFPs in the theta frequency band for fixations on targets compared to fixations on distractors (**Fig. 3**). Specifically, foveal units demonstrated target-induced desynchronization between V4 spike and V4 LFP (**Fig. 3A**; two-tailed two-sample *t*-test: *t*(58810) = 20.53, P = 2.60×10^−93^; Bonferroni correction for multiple comparisons), between V4 spike and IT LFP (**Fig. 3B**; *t*(15604) = 7.48, P = 7.55×10^−14^), between IT spike and V4 LFP (**Fig. 3E**; *t*(15960) = 13.98, P = 3.54×10^−44^), between IT spike and IT LFP (**Fig. 3F**; *t*(45318) = 14.48, P = 1.99×10^−47^), between IT spike and OFC LFP (**Fig. 3G**; *t*(10076) = 5.00, P = 5.77×10^−7^), and between OFC spike and OFC LFP (**Fig. 3J**; *t*(14068) = 3.87, P = 1.10×10^−4^). Peripheral units only demonstrated target-induced desynchronization between V4 spike and V4 LFP (**Fig. 3A**; *t*(16674) = 8.27, P = 1.43×10^−16^), and between LPFC spike and LPFC LFP (**Fig. 3L**; *t*(10418) = 4.80, P = 1.62×10^−6^). Therefore, the coherence of foveal units was more extensively modulated by attention for target response than the coherence of peripheral units.

**Fig. 3.**
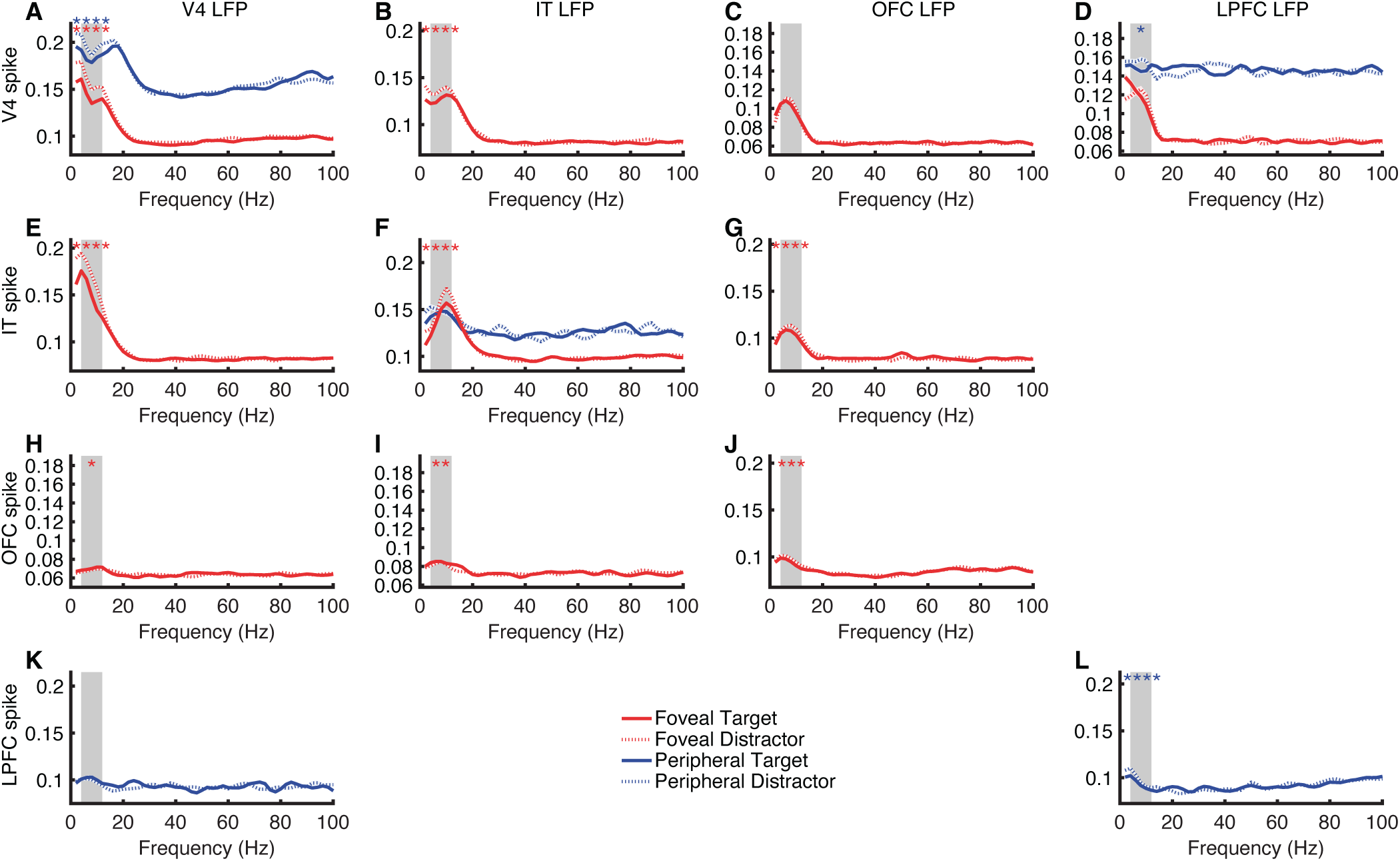
Spike-LFP coherence for feature attention effect. **(A)** V4 spike-V4 LFP. **(B)** V4 spike-IT LFP. **(C)** V4 spike-OFC LFP. **(D)** V4 spike-LPFC LFP. **(E)** IT spike-V4 LFP. **(F)** IT spike-IT LFP. **(G)** IT spike-OFC LFP. **(H)** OFC spike-V4 LFP. **(I)** OFC spike-IT LFP. **(J)** OFC spike-OFC LFP. **(K)** LPFC spike-V4 LFP. **(L)** LPFC spike-LPFC LFP. Red: coherence between units with a focal foveal receptive field (RF). Blue: coherence between units with overlapping peripheral RFs. Solid line: target in RF. Dotted line: distractor in RF. Red and blue shaded areas denote ±SEM across spike-LFP pairs. Gray shaded area denotes the theta frequency band (4 - 12 Hz). Asterisks indicate a significant difference in target-induced desynchronization (i.e., the reduction in spike-LFP coherence for target in RF compared to distractor in RF, averaged across the theta frequency band) using two-tailed two-sample *t*-test (uncorrected). *: P < 0.05, **: P < 0.01, ***: P < 0.001, and ****: P < 0.0001. Red: foveal units. Blue: peripheral units.

We next investigated the modulation by category selectivity by comparing the difference in coherence between faces versus houses. Foveal units (regardless of category selectivity based on firing rate) exhibited substantially stronger spike-LFP coherence in the theta frequency band for faces compared to houses across brain areas. This included coherence between V4 spike and IT LFP (**Fig. 4B**; two-tailed two-sample *t*-test: *t*(15604) = 18.98, P = 1.89×10^−79^; Bonferroni correction for multiple comparisons), between V4 spike and OFC LFP (**Fig. 4C**; *t*(9002) = 4.66, P = 3.18×10^−6^), between IT spike and V4 LFP (**Fig. 4E**; *t*(15960) = 37.64, P = 4.00×10^−297^), between IT spike and IT LFP (**Fig. 4F**; *t*(45318) = 51.86, P < 10^−50^), between IT spike and OFC LFP (**Fig. 4G**; *t*(10076) = 10.16, P = 3.98×10^−24^), between OFC spike and V4 LFP (**Fig. 4H**; *t*(8312) = 11.43, P = 4.95×10^−30^), between OFC spike and IT LFP (**Fig. 4I**; *t*(9354) = 23.45, P = 3.50×10^−118^), and between OFC spike and OFC LFP (**Fig. 4J**; *t*(14068) = 9.17, P = 5.52×10^−20^). However, peripheral units only exhibited a significant difference in coherence between V4 spike and LPFC LFP (**Fig. 4D**; *t*(3726) = 3.67, P = 2.45×10^−4^), with houses having a stronger coherence than faces. Therefore, the coherence of foveal units was more extensively modulated by category selectivity than the coherence of peripheral units.

**Fig. 4.**
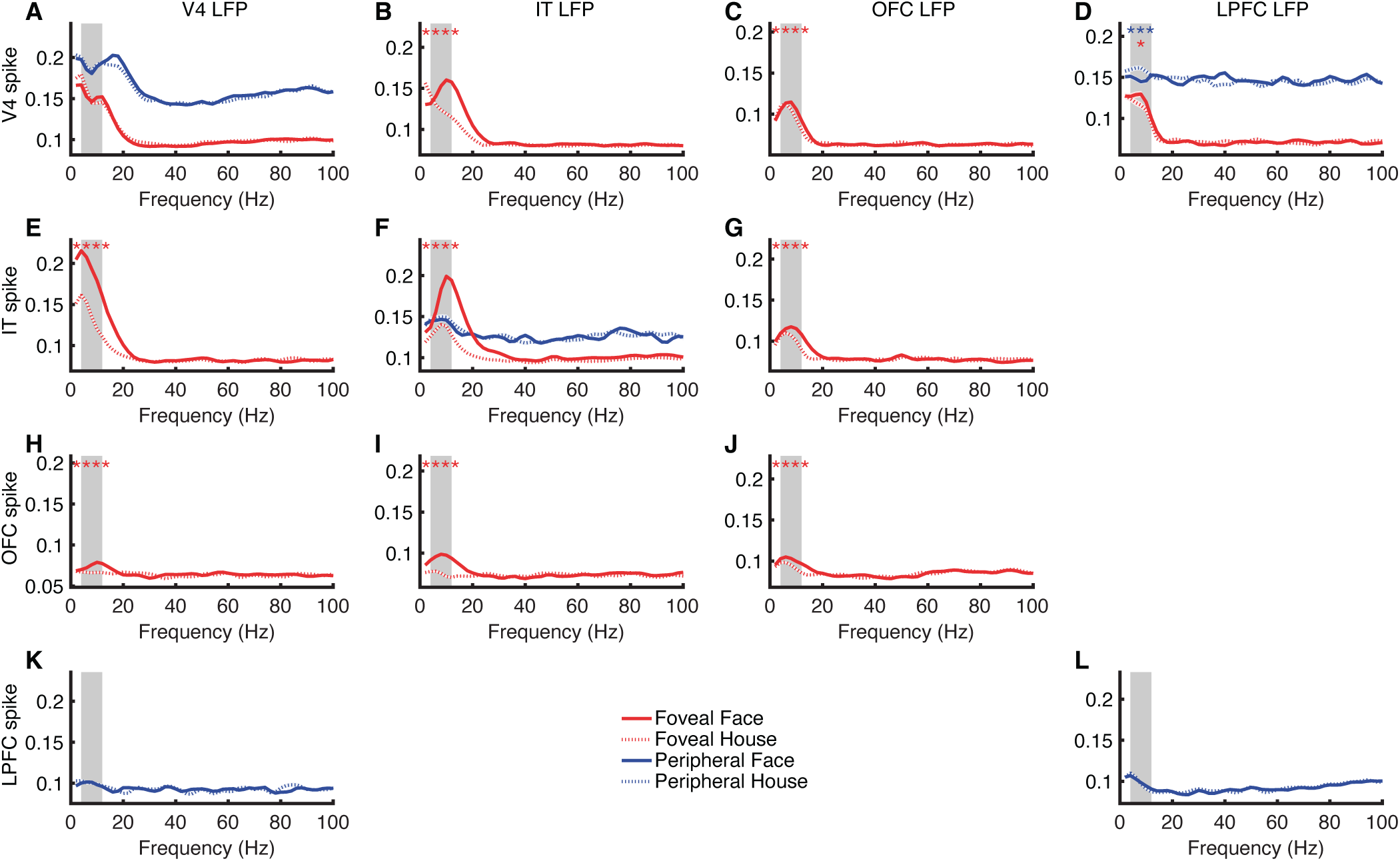
Spike-LFP coherence for category selectivity. **(A)** V4 spike-V4 LFP. **(B)** V4 spike-IT LFP. **(C)** V4 spike-OFC LFP. **(D)** V4 spike-LPFC LFP. **(E)** IT spike-V4 LFP. **(F)** IT spike-IT LFP. **(G)** IT spike-OFC LFP. **(H)** OFC spike-V4 LFP. **(I)** OFC spike-IT LFP. **(J)** OFC spike-OFC LFP. **(K)** LPFC spike-V4 LFP. **(L)** LPFC spike-LPFC LFP. Red: coherence between units with a focal foveal receptive field (RF). Blue: coherence between units with overlapping peripheral RFs. Solid line: face in RF. Dotted line: house in RF. Red and blue shaded areas denote ±SEM across spike-LFP pairs. Gray shaded area denotes the theta frequency band (4 - 12 Hz). Asterisks indicate a significant difference in synchronization (i.e., the difference in spike-LFP coherence for face in RF compared to house in RF, averaged across the theta frequency band) using two-tailed two-sample *t*-test (uncorrected). *: P < 0.05, **: P < 0.01, ***: P < 0.001, and ****: P < 0.0001. Red: foveal units. Blue: peripheral units.

Together, foveal units differentially engaged the attention and visual coding network compared to peripheral units.

### Coordination between foveal and peripheral processing for spatial attention

We next investigated the interaction and coordination between foveal and peripheral units/LFPs when encoding spatial attention and saccade selection.

First, we observed that target-induced desynchronization (i.e., reduced spike-LFP coherence for targets in RF compared to distractors in RF) was only present for peripheral units when the fixations were on distractors (**Fig. 5A**; two-tailed two-sample *t*-test: *t*(16228) = 3.38, P = 7.24×10^−4^; Bonferroni correction for multiple comparisons) but not targets (**Fig. 5B**; P > 0.05). This result suggested that the processing in the fovea influenced the processing in the periphery, consistent with our prior report [16].

**Fig. 5.**
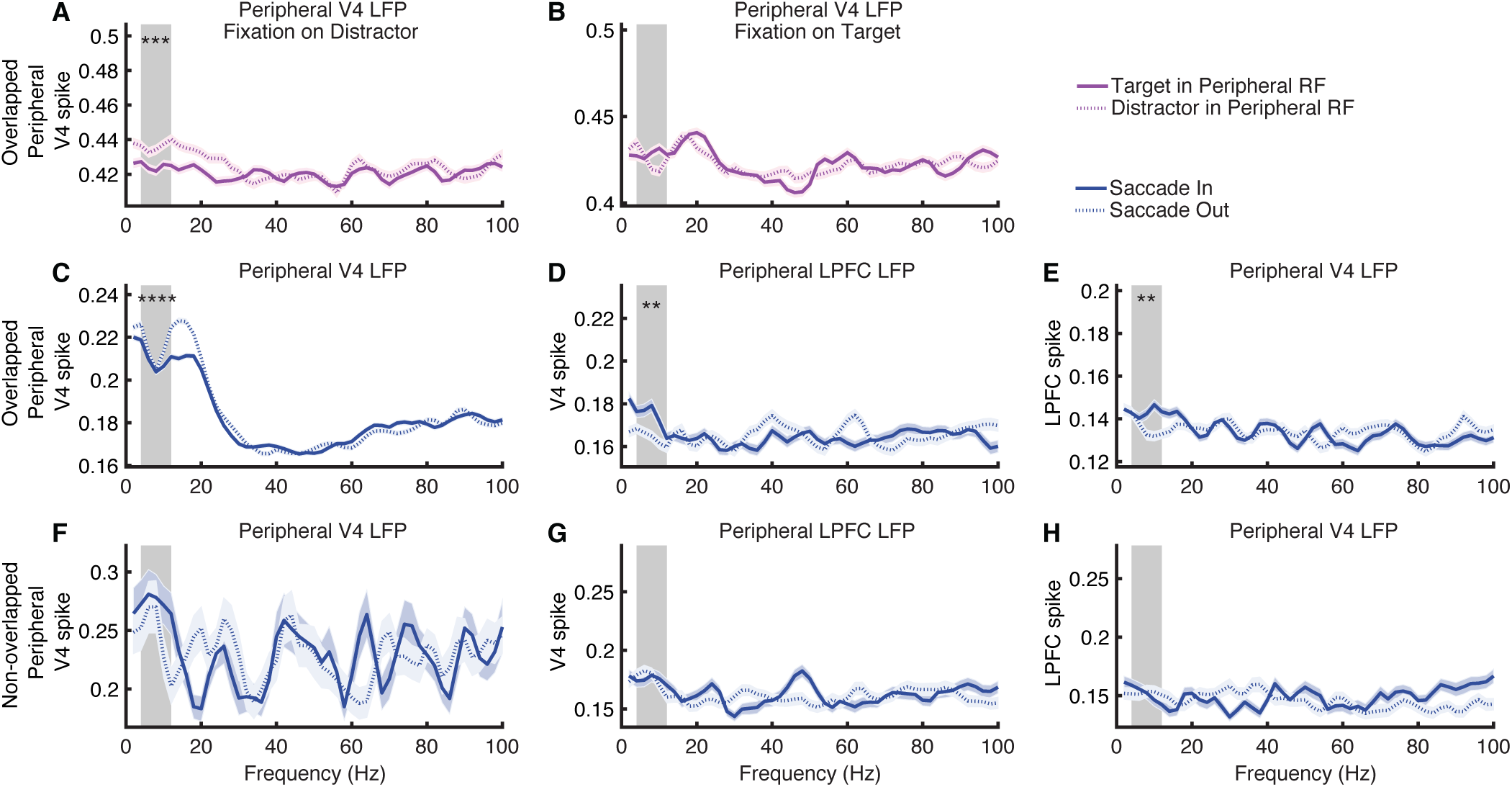
Spike-LFP coherence for spatial attention. **(A, B)** V4 spike-V4 LFP coherence between units and LFPs that had overlapping peripheral receptive fields (RFs). **(A)** Foveal fixation on distractors. **(B)** Foveal fixation on targets. Solid line: target in the peripheral RF. Dotted line: distractor in the peripheral RF. **(C-E)** Coherence between units and LFPs that had overlapping peripheral RFs. **(F-H)** Coherence between units and LFPs that had non-overlapping peripheral RFs. Solid line: saccade in (i.e., the next fixation was within the RF). Dotted line: saccade out (i.e., the next fixation was not within the RF). **(C, F)** V4 spike-V4 LFP. **(D, G)** V4 spike-LPFC LFP. **(E, H)** LPFC spike-V4 LFP. Magenta and blue shaded areas denote ±SEM across spike-LFP pairs. Gray shaded area denotes the theta frequency band (4 - 12 Hz). Asterisks indicate a significant difference in synchronization using two-tailed two-sample *t*-test (uncorrected). *: P < 0.05, **: P < 0.01, ***: P < 0.001, and ****: P < 0.0001.

Second, saccading to a peripheral RF (i.e., “saccade in”) elicited a stronger coherence in the theta frequency band between V4 spike and LPFC LFP (**Fig. 5D**; two-tailed two-sample *t*-test: *t*(2900) = 2.94, P = 0.003; Bonferroni correction for multiple comparisons) and between LPFC spike and V4 LFP (**Fig. 5E**; *t*(2468) = 2.72, P = 0.007) compared to saccading outside of a peripheral RF (i.e., “saccade out”). In contrast, saccade-in conditions had a reduced coherence between V4 spike and V4 LFP compared to saccade-out conditions (**Fig. 5C**; *t*(16474) = 4.19, P = 2.79×10^−5^). Therefore, the coherence of peripheral units could predict whether the next fixation landed in the peripheral RF. Notably, these differences between saccade-in versus saccade-out conditions were only present when spikes and LPFs shared overlapping RFs (**Fig. 5C-E** versus **Fig. 5F-H**; all Ps > 0.05 for **Fig. 5F-H**), suggesting that the processing of saccade selection was specific to RFs.

Lastly, we analyzed the interaction and coordination between foveal and peripheral processing for spatial attention and saccade selection. We found that during fixations on distractors, the coherence between peripheral V4 spike and foveal IT LFP (**Fig. 6B**; two-tailed two-sample *t*-test: *t*(8598) = 7.87, P = 3.99×10^−15^; Bonferroni correction for multiple comparisons), the coherence between peripheral V4 spike and foveal OFC LFP (**Fig. 6C**; *t*(7100) = 7.63, P = 2.57×10^−14^), and the coherence between peripheral IT spike and foveal OFC LFP (**Fig. 6F**; *t*(1350) = 3.10, P = 0.002) could predict saccade selection (i.e., “saccade in” versus “saccade out”). Furthermore, during fixations on targets, the coherence between peripheral V4 spike and foveal IT LFP (**Fig. 6B**; *t*(8270) = 4.11, P = 3.98×10^−5^) could predict saccade selection. Therefore, the interaction and coordination between foveal and peripheral processing were engaged in spatial attention and saccade selection. Notably, such coordination was primarily through peripheral spikes and foveal LFP: the coherence between foveal spike and peripheral LFP (**Fig. S3**) could barely predict saccade selection. Furthermore, the coherence between spikes from units with a broad foveal RF and peripheral LFP could barely predict saccade selection either (**Fig. S4**). Therefore, our results indicated specificity and direction of coordination between foveal and peripheral processing.

**Fig. 6.**
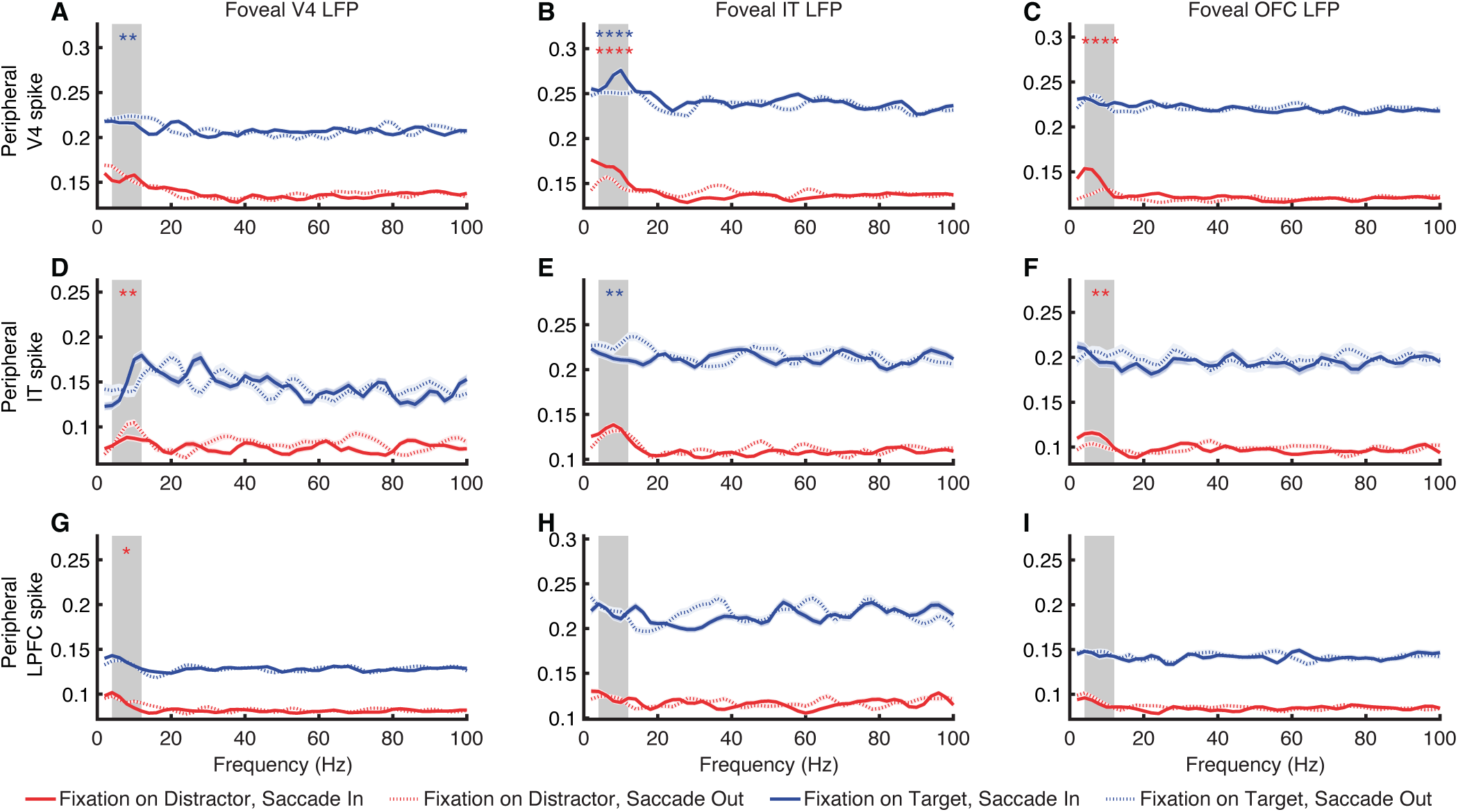
Spike-LFP coherence between peripheral spikes and foveal LFPs for spatial attention. **(A)** V4 spike-V4 LFP. **(B)** V4 spike-IT LFP. **(C)** V4 spike-OFC LFP. **(D)** IT spike-V4 LFP. **(E)** IT spike-IT LFP. **(F)** IT spike-OFC LFP. **(G)** LPFC spike-V4 LFP. **(H)** LPFC spike-IT LFP. **(I)** LPFC spike-OFC LFP. Red line: current foveal fixation on distractors. Blue line: current foveal fixation on targets. Solid line: saccade in (i.e., the next fixation was within the RF). Dotted line: saccade out (i.e., the next fixation was not within the RF). Red and blue shaded areas denote ±SEM across spike-LFP pairs. Gray shaded area denotes the theta frequency band (4 - 12 Hz). Asterisks indicate a significant difference in synchronization using two-tailed two-sample *t*-test (uncorrected). *: P < 0.05, **: P < 0.01, ***: P < 0.001, and ****: P < 0.0001.

Together, our results suggest interaction and coordination between foveal and peripheral processing for spatial attention and saccade selection.

### Relationship between behavior and spike-LFP coherence

Finally, we investigated the relationship between search efficiency (indexed by RT and the number of fixations) and target-induced desynchronization (indexed by the reduction in spike-LFP coherence for targets compared to distractors) in foveal and peripheral units. Specifically, we found that RT significantly correlated with the target-induced reduction in IT spike-V4 LFP coherence (**Fig. 7A**; Pearson correlation: *r* = −0.18, P = 7.62×10^−5^; Bonferroni correction for multiple comparisons), V4 spike-V4 LFP coherence (**Fig. 7B**; *r* = −0.08, P = 9.81×10^−4^), and IT spike-IT LFP coherence (**Fig. 7C**; *r* = −0.18, P = 2.92×10^−11^), in foveal units. Target-induced reduction in V4 spike-IT LFP coherence in peripheral units (**Fig. 7E**; *r* = −0.41, P = 1.45×10^−5^) could also predict RT, and interestingly, V4 spike-V4 LFP coherence in peripheral units exhibited a positive correlation (**Fig. 7D**; *r* = 0.16, P = 6.79×10^−6^), indicating that target-induced reduction in coherence within V4 negatively impacted search efficiency. Similarly, the number of fixations per trial significantly correlated with the target-induced reduction in IT spike-V4 LFP coherence (**Fig. 7F**; *r* = −0.19, P = 2.60×10^−5^), V4 spike-V4 LFP coherence (**Fig. 7G**; *r* = −0.07, P = 3.01×10^−3^), and IT spike-IT LFP coherence (**Fig. 7H**; *r* = −0.18, P = 7.15×10^−11^) in foveal units, as well as V4 spike-V4 LFP coherence (**Fig. 7I**; *r* = 0.18, P = 3.25×10^−7^; note the positive correlation) and V4 spike-IT LFP coherence (**Fig. 7J**; *r* = −0.38, P = 6.52×10^−5^) in peripheral units. However, we did not observe significant correlations between search efficiency and feature attention effects (all Ps > 0.05), nor between feature attention effects and target-induced desynchronization (all Ps > 0.05).

**Fig. 7.**
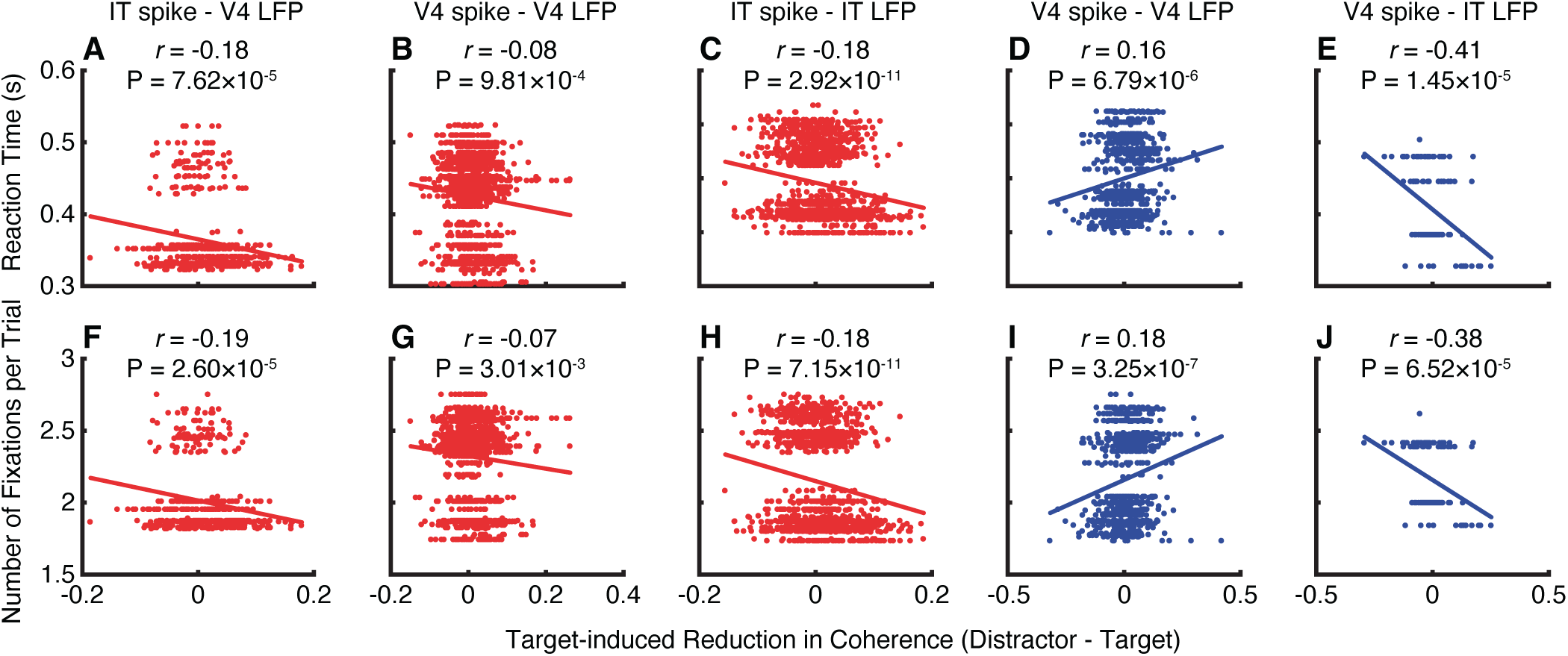
Relationship between behavior and spike-LFP coherence. **(A-E)** Correlation between reaction time (RT) and target-induced reduction in spike-LFP coherence (distractor − target). **(F-J)** Correlation between the number of fixations and target-induced reduction in spike-LFP coherence (distractor − target). **(A, F)** IT spike-V4 LFP coherence for foveal units. **(B, G)** V4 spike-V4 LFP coherence for foveal units. **(C, H)** IT spike-IT LFP coherence for foveal units. **(D, I)** V4 spike-V4 LFP coherence for peripheral units. **(E, J)** V4 spike-IT LFP coherence for peripheral units. Each dot represents a unit, and the lines represent the linear fit. Red: foveal units. Blue: peripheral units.

Together, we showed that target-induced desynchronization within the temporal cortex could explain search efficiency: the search became more efficient (reduced RT and number of fixations) with increasing desynchronization. Notably, foveal and peripheral units engaged different neural substrates and exhibited different correlations between neural responses and search behavior, further indicating that they differentially engaged the attention network.

## Discussion

In this study, we simultaneously recorded from a large number of units in area V4, IT cortex, OFC, and LPFC while monkeys performed a free-gaze visual search task. In each brain area, we identified foveal and peripheral units and the proportion of foveal versus peripheral units varied substantially across brain areas. Foveal and peripheral units exhibited systematic differences in encoding visual information and visual attention. At the neural circuit and network level, we found that the spike-LFP coherence of foveal units was more extensively modulated by both feature attention and visual selectivity. Therefore, foveal units differentially engaged the attention and visual coding network compared to peripheral units. Furthermore, we delineated the interaction and coordination between foveal and peripheral processing for spatial attention and saccade selection. Finally, we showed that the search became more efficient with increasing target-induced desynchronization within the temporal cortex, and foveal and peripheral units engaged different neural substrates and exhibited different correlations between neural responses and search behavior. Together, our results have provided important insights into how the brain processes and integrates visual information for active visual behaviors.

Our previous study revealed a novel distribution of feature attention effects across the entire visual field [16]. In the present study, we further revealed systematic differences in response profile and functional connectivity between foveal units and peripheral units. Moreover, our prior study revealed a bi-directionally interactive loop between parallel feature attention and serial saccade selection [16]. Our present study further corroborated these findings by showing the interaction and coordination between foveal and peripheral processing for spatial attention and saccade selection. In a recent study investigating the RFs of visual neurons during natural vision of a large number of images in primates [48], it has been shown that feature-selective responses remain largely eye-centered, with neurons tracking their RF contents as the eyes move. This suggests a predominantly gaze-dependent reference frame for visual neurons during natural vision, with limited evidence for predictive remapping or integration based on viewing history.

In monkeys, studies of visual coding have primarily focused on the central visual field [49–54]. Conversely, research on spatial attention and feature attention has predominantly explored the peripheral visual field [8, 10]. In this study, we trained monkeys to perform a free-gaze visual search task using natural face and object stimuli, enabling detailed analysis of both visual attention and visual selectivity across the entire visual field. On the one hand, we observed not only target-selective units modulated by visual attention (**Fig. 2**) but also desynchronization for attended stimuli (fixations on search targets) in the theta frequency band compared to the same stimuli when they were unattended (fixations on distractors) (**Fig. 3**). Notably, this target-induced desynchronization primarily appeared in foveal units (**Fig. 3**). Previous research has shown desynchronization for attended stimuli in V4 within a similar frequency band [34, 36]. Desynchronization in V4 spike-V4 LFP coherence, V4 spike-frontal eye field (FEF) LFP coherence, and FEF spike-V4 LFP coherence has been observed for feature-based attention, where discrimination between target and distractor in the peripheral RF occurs, as well as during saccade selection, involving directing attention into (i.e., saccade in) or out of (i.e., saccade out) the peripheral RF [36]. On the other hand, across brain areas, we observed above-chance populations of category-selective units that differentiated between faces and houses, notably only in foveal units (**Fig. 2**). We also observed that foveal units exhibited substantially stronger spike-LFP coherence in the theta frequency band for faces compared to houses across brain areas (**Fig. 4**), and importantly, the coherence of foveal units was more extensively modulated by category selectivity than the coherence of peripheral units (**Fig. 4**).

While numerous studies have illustrated visual category selectivity along the ventral processing stream [55–57] (see [50, 53, 54] for more recent analyses of visual features in IT), our study is among the first to examine the interaction between visual attention and visual selectivity (**Fig. 2A-C**). Specifically, we observed that the proportion of units qualifying as both category-selective and target-selective was not greater than expected from independence of these two attributes, suggesting that visual coding and attention coding remained independent in V4, IT, and OFC. Like our current findings, neurons in the human amygdala and hippocampus not only encode visual attention directed towards visual search targets but also encode visual categories [58]. Importantly, the neurons encoding visual attention and visual categories are largely from separate populations. It is also worth noting that in humans, single-neuron correlates of visual attention during visual search have primarily been investigated using the central visual field [58, 59]. Although target-selective and category-selective units were selected during different periods of the task, and cells (especially prefrontal cells) may be subject to context modulation, it is worth noting that we did not directly compare these two groups of units. Instead, we aimed to minimize interactions between the two groups of units by ensuring there was a single stimulus in the RF when studying category selectivity, and by matching the stimulus between fixations on targets and distractors when studying the feature attention effect. Future studies will be needed to further investigate the context modulation of these units.

Spatial attention and feature attention have along be regarded as different processes [8]. Spatial attentional modulation of neural response mainly depends on the relationship between the location of the stimulus in the RF and the attended spatial location, while feature attention can simultaneously act on multiple visual stimuli across the entire visual field [33]. Feature attention has a parallel distributed global effect, independent of the next saccade location, and this global feature attention can guide subsequent spatial attention in the visual search process [8]. In visual search, there are parallel processing and serial selection processes, benefiting from the modulation of feature and spatial attention, respectively. Spatial attention effects occur serially with the continuous change of saccadic targets, while feature attention effects occur in parallel [8, 60]. In this study, we investigated the functional connectivity for units encoding spatial and feature attention and demonstrated systematic differences between foveal and peripheral units for both attention processes.

We found that the OFC and LPFC within the prefrontal cortex (PFC), along with their interactions with the temporal cortex (V4 and IT), played an important role in visual attention. Functional magnetic resonance imaging (fMRI) studies in humans have shown that a network including the PFC may generate intrinsic attention signals [61]. Studies in non-human primates have provided neurophysiological evidence for the sources of attentional control, which include the ventral prearcuate area (VPA) and frontal eye field (FEF) regions of the PFC [12]. Neural activity induced by attention first appears in the PFC, approximately 30-50 milliseconds before it appears in area V4, suggesting top-down modulation of attention [12, 62]. Supporting this notion, lesion of the LPFC in macaques weakens the modulatory effects of attention on the visual cortex [63]. Furthermore, deactivation of the VPA reduces feature attention effects in V4 and impairs performance in behavioral tasks [12, 14], suggesting that the VPA is the source of top-down signals for feature-based attentional control, which facilitate the processing of relevant stimuli across the entire visual field. In addition, control signals for feature-based attention also originate from the posterior IT cortex. Electrical stimulation of this area has been shown to alter the allocation of attention [64]. Lastly, attention can modulate the representation of stimulus value in OFC neurons [43, 44]. Given the substantial difference in the proportion of foveal versus peripheral units across the PFC (**Fig. 1B**), as well as systematic differences in response profile (**Fig. 2**) and functional connectivity (**Fig. 3** and **Fig. 4**) between foveal and peripheral units, future studies are needed to investigate the role of the PFC in visual attention relative to RF location.

## Methods

### Subjects

Two male rhesus macaques, weighing 12 and 15 kg, were used in the study. The monkeys were implanted under aseptic conditions with a post to fix the head and recording chambers over areas V4, IT (including both TE and TEO), LPFC, and OFC. The localization of the chambers was based on MRI scans obtained before surgery. All experiments were performed at the Shenzhen Institutes of Advanced Technology, Chinese Academy of Sciences, with the approval of the Institutional Animal Care and Use Committee (No. SIAT-IRB-160223-NS-ZHH-A0187-003). This dataset has been analyzed in previous studies [16].

### Tasks and stimuli

Monkeys were trained to perform a free-gaze visual search task. A central fixation was presented for 400 ms, followed by a cue lasting 500 to 1300 ms. After a delay of 500 ms, the search array was on. The search array contained 11 items, including two targets, randomly selected from a total of 20 predefined locations. Monkeys were required to find either one of the two targets within 4000 ms and maintain fixation on the target for 800 ms to receive a juice reward. No constraints were placed on their search behavior to allow animals to perform the search naturally. Before the onset of the search array, monkeys were required to maintain a central fixation. The two target stimuli belonged to the same category as the cue stimulus, though they were distinct images. We utilized four categories of stimuli—face, house, flower, and hand—each comprising 40 images. The cue stimulus was randomly selected from the house or face stimuli with equal probability. The remaining 9 stimuli in the search array were drawn from the other three categories. Each stimulus subtended an area of approximately 2° × 2°, with the hue, saturation in the HSV color space, aspect ratio, and luminance of these images matched across categories. The 20 locations, covering the visual field of eccentricities from 5° to 11°, included 18 locations located symmetrically in the left and right visual field, with 9 on each side, and 2 locations on the vertical middle line (**Fig. 1D**).

A visually guided saccade task was employed to map the peripheral receptive fields (RFs) of recorded units. Following a 400-ms central fixation, a stimulus (face or house; the same stimuli as the visual search task) randomly appeared in one of the 20 locations, and monkeys were required to make a saccade to the stimulus within 500 ms and maintain fixation on it for 300 ms to receive a reward. Behavioral experiments were conducted using the MonkeyLogic software (University of Chicago, IL), which presented the stimuli, monitored eye movements, and triggered the delivery of the reward.

### Electrophysiology

Single-unit and multi-unit spikes were recorded from V4, IT, LPFC, and OFC using 24- or 32-contact electrodes (V-Probe or S-Probe, Plexon Inc, Dallas, USA) in a 128-channel Cerebus System (Blackrock Microsystems, Salt Lake City, UT, USA). In most sessions, we recorded activities in two of the areas simultaneously. Neural recordings were filtered between 250 Hz and 5 kHz and digitized at 30 kHz to obtain spike data. Spike sorting was performed using Plexon’s Offline Sorter™ (OFS). Neural recordings were filtered between 0.3 and 250 Hz and digitized at 1000 Hz to obtain LFP signals. The recording locations in V4, IT, LPFC, and OFC were verified with MRI. Eye movements were recorded using an infrared eye-tracking system (iViewX Hi-Speed, SensoMotoric Instruments (SMI), Teltow, Germany) at a sampling rate of 500 Hz.

### Data analysis: spike rate

Measurements of neural activity were obtained from spike density functions, which were generated by convolving the time of action potentials with a function that projects activity forward in time (Growth = 1 ms, Decay = 20 ms) and approximates an EPSP [65]. Specifically, this spike density function has two advantages [65]. First, each spike exerts influence only forward in time, representing the actual postsynaptic effect of each cell. Second, by using a function that resembles a postsynaptic potential, we can apply time constants similar to those measured physiologically. Additionally, using the postsynaptic potential filter for time course analysis is advantageous because, when using the Gaussian filter, target discrimination times sometimes occur earlier than the neuron’s evident visual latency. This impossible outcome occurs because, with the Gaussian filter, spikes exert influence backward in time. The spike rate of each unit was normalized by the mean baseline firing rate during the fixation spot preceding the cue.

### Data analysis: receptive field

The visual response to the cue and the search array in the free-gaze visual search task was assessed by comparing the firing rate during the post-stimulus period (50 to 200 ms after cue/array onset) to the corresponding baseline (−150 to 0 ms relative to cue/array onset) using a Wilcoxon rank-sum test. Based on these responses, we classified units into three categories of RFs:

i. Units with a focal foveal RF: These units responded solely to the cue in the foveal region (P < 0.05) but not to the search array that included items in the periphery (P > 0.05).
ii. Units with a broad foveal RF: These units responded to both the cue and the search array (both Ps < 0.05).
iii. Units with a peripheral RF: These units only responded to the search array (P < 0.05) but not to the cue (P > 0.05). The RFs of these units were additionally mapped based on their activities in the visually guided saccade task. Units whose RFs could be mapped in this task had a localized peripheral RF, whereas units whose RFs could not be mapped had an unlocalized peripheral RF (i.e., units that responded to the search array onset but not during the saccade task; 11.87% of visually responsive units in V4, 5.58% in IT, 3.20% in OFC, and 58.29% in LPFC had an unlocalized RF).

Units not classified into the above categories (both Ps > 0.05) were not visually responsive and were excluded from further analysis.

### Data analysis: category selectivity

We determined the category selectivity of each unit by comparing the response to face cues versus house cues in a time window of 50 to 200 ms after cue onset (Wilcoxon rank-sum test, P < 0.05). We further imposed a second criterion using a selectivity index similar to indices employed in previous IT studies [66, 67]. For each unit with a foveal RF, the response to face stimuli (*R*_face_) or house stimuli (*R*_house_) was calculated using the visual search task by subtracting the mean baseline activity (−150 to 0 ms relative to the onset of the cue) from the mean response to the face or house cue (50 to 200 ms after the onset of the cue). For each unit with a peripheral RF, *R*_face_ and *R*_house_ were calculated using the visually guided saccade task by subtracting the mean baseline activity (−150 to 0 ms relative to the peripheral stimulus onset) from the mean response to the saccade target (50 to 200 ms after the onset of the saccade target). It is worth noting that for both foveal and peripheral units, we ensured that there was only one stimulus in the RF. The selectivity index (SI) was then defined as (*R*_face_ − *R*_house_) / (*R*_face_ + *R*_house_). SI was set to 1 when *R*_face_ > 0 and *R*_house_ < 0, and to −1 when *R*_face_ < 0 and *R*_house_ > 0. Face-selective units were required to have an *R*_face_ at least 130% of *R*_house_ (i.e., the corresponding SI was greater than 0.13). Similarly, house-selective units were required to have an *R*_house_ at least 130% of *R*_face_ (i.e., the corresponding SI was smaller than −0.13). Units were labeled as non-category-selective if the response to face cues versus house cues was not significantly different (P > 0.05). The remaining units that did not fit into any of the aforementioned types were classified as undefined units (i.e., there was a significant difference but did not meet the second criterion). It is worth noting that we did not use the activity during the search to calculate the SI to minimize interactions with the feature attention effect (see below) and between RFs.

### Data analysis: selection of target-selective units

We used the mean firing rate in a time window of 150 ms to 225 ms after fixation onset as the response to each fixation. For each unit, if there was a significant difference in response (determined using a two-tailed Wilcoxon signed-rank test, with a significance threshold of P < 0.05) between fixations on targets and fixations on distractors, it was classified as an *target-selective unit*. Similarly, for units with a peripheral RF (as described above), we compared the response between targets and distractors within the RF in the same time window as for foveal units. Lastly, we calculated the feature attention effect as the difference in firing rate between the same stimuli when they served as targets versus distractors.

### Data analysis: differential latency

For each unit, the latency was defined as the first 20-ms bin out of twelve successive bins that had a significantly different response between conditions (i.e., preferred versus non-preferred, or targets versus distractors) using a two-tailed Wilcoxon signed-rank test with a significance threshold of P < 0.05.

### Data analysis: spike-LFP coherence

We implemented the spike-LFP coherence analysis using the Chronux toolbox (www.chronux.org) in MATLAB. LFP signals were preprocessed by removing the power-line artifact using a method described in a previous study [68]. Specifically, for each LFP epoch of interest, we took a 10 s epoch from the continuous signal with the epoch of interest in the middle, filtered it at 1 to 100 Hz, and calculated the discrete Fourier transform (DFT) of the 10 s epoch at 50 Hz without any tapering. We then constructed a 50 Hz sine wave with the amplitude and phase as estimated by the DFT and subtracted this sine wave (i.e., the estimated power-line artifact) from the 10 s epoch. We used a single Hanning taper across frequencies, but we derived similar results using multitaper methods for higher frequencies (> 25 Hz) [68]. Coherence between two signals, *x* and *y*, was calculated using the following formula:

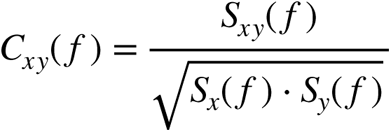

where *S_x_*(*f*) and *S_y_*(*f*) denote the auto-spectra and *S_xy_*(*f*) represents the cross-spectrum of the two signals *x* and *y*. Auto-spectra and cross-spectra were averaged across trials before coherence calculation. We used a 200-ms time window for each fixation (0 to 200 ms relative to fixation onset). We chose this time window given the mean fixation duration (208 ms). Furthermore, it elicited stronger coherence than other windows (note that spike-LFP coherence can represent a different neural mechanism compared to the firing rate). Notably, we used an equal number of fixations and an equal number of spikes between conditions to calculate coherence for a given pair of recording sites, thus eliminating bias from different sample sizes.

We used spikes from all channels, but based the analysis on the selectivity of the units (i.e., category selectivity, attention selectivity, and RFs). To avoid spikes contributing to the LFP recorded on the same channel, we excluded the LFP channel from which the spike originated. In other words, we always utilized signals from two different channels to calculate coherence. For the analysis of category selectivity and feature attention effect, we did not select LFPs based on their category selectivity, attention selectivity, or the selectivity of the associated units (e.g., the LFP signals could come from channels with both face-selective units and house-selective units). For the analysis with matching RFs, we defined the RF of an LFP channel as the combined RF of all units from that channel and selected LFPs to match the RF of the corresponding units.

## Acknowledgements

This research was supported by the NSF (BCS-1945230), NIH (R01MH129426), and AFOSR (FA9550-21-1-0088). The funders had no role in study design, data collection and analysis, decision to publish, or preparation of the manuscript.

## Author Contributions

J.Z. and H.Z. designed research. J.Z. and X.Z. performed experiments. J.Z. and S.W. analyzed data. J.Z., H.Z., and S.W. wrote the paper. All authors discussed the results and contributed toward the manuscript.

---

**Fig. S1.**
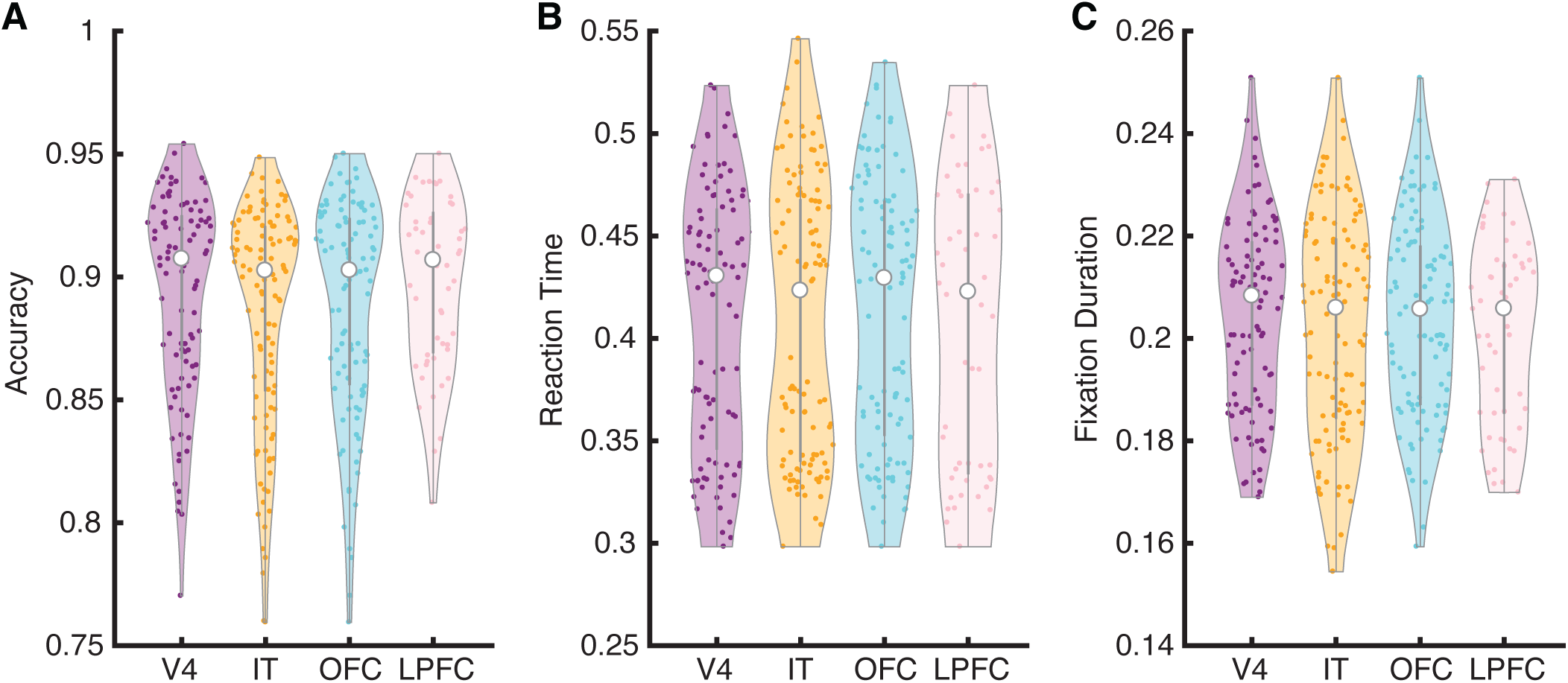
Behavior for sessions with recordings from different brain regions. **(A)** Search accuracy. **(B)** Reaction time. **(C)** Fixation Duration. In the violin plots, the white dot represents the median, the thick gray bar in the center represents the interquartile range, the thin gray line represents the rest of the distribution, except for points that are determined to be outliers using a method that is a function of the interquartile range. On each side of the gray line is a kernel density estimation to show the distribution shape of the data. Each dot represents an individual recording session.

**Fig. S2.**
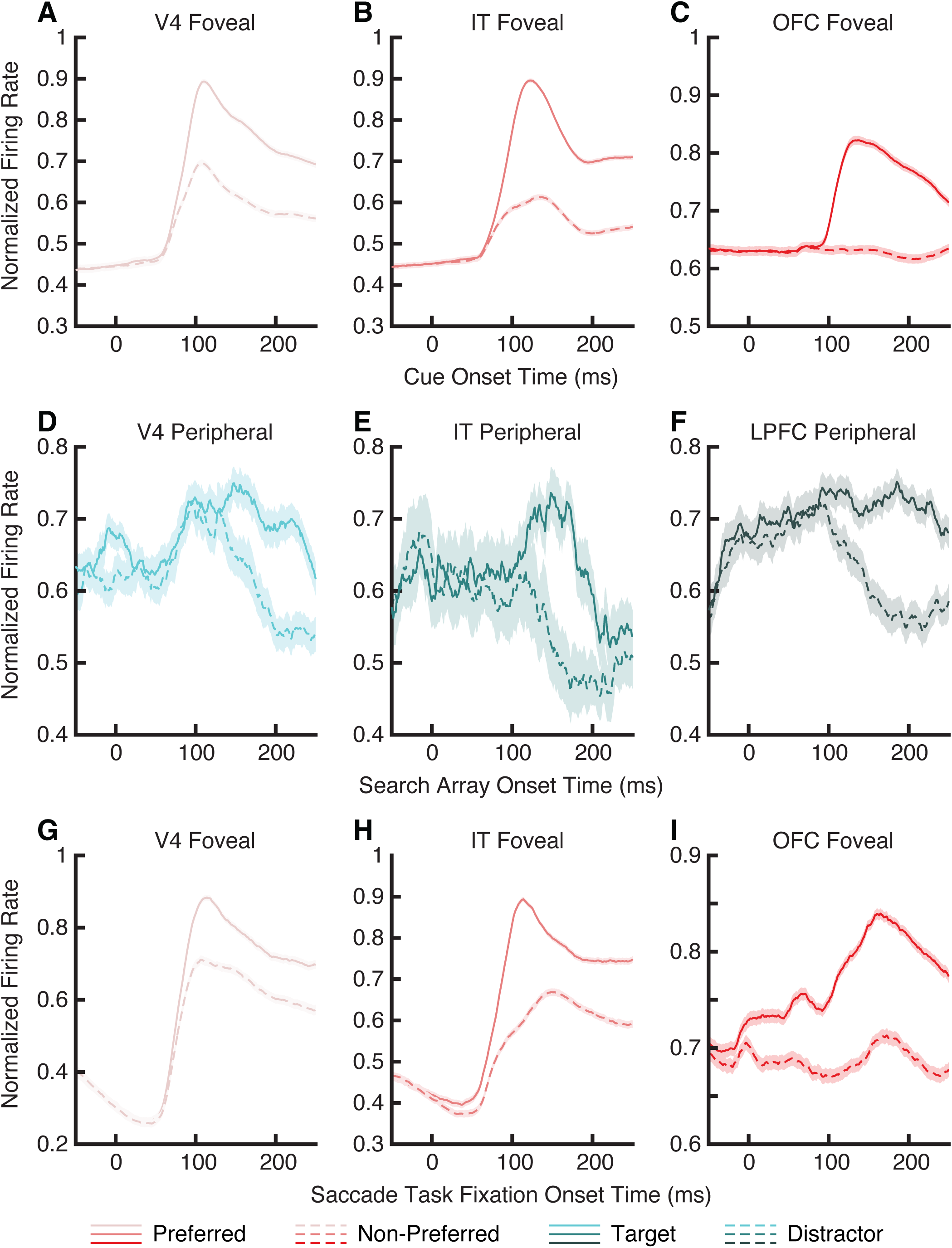
Neural response time-locked at different events. **(A-C)** Category selectivity of foveal units when time-locked at cue onset. **(D-F)** Feature attention effect of peripheral units when time-locked at search array onset. **(G-I)** Category selectivity of foveal units after foveation of the saccade target in the visually guided saccade task. Legend conventions as in Fig. 2. In these plots, we normalized the firing rate of each unit by its maximum rate across conditions.

**Fig. S3.**
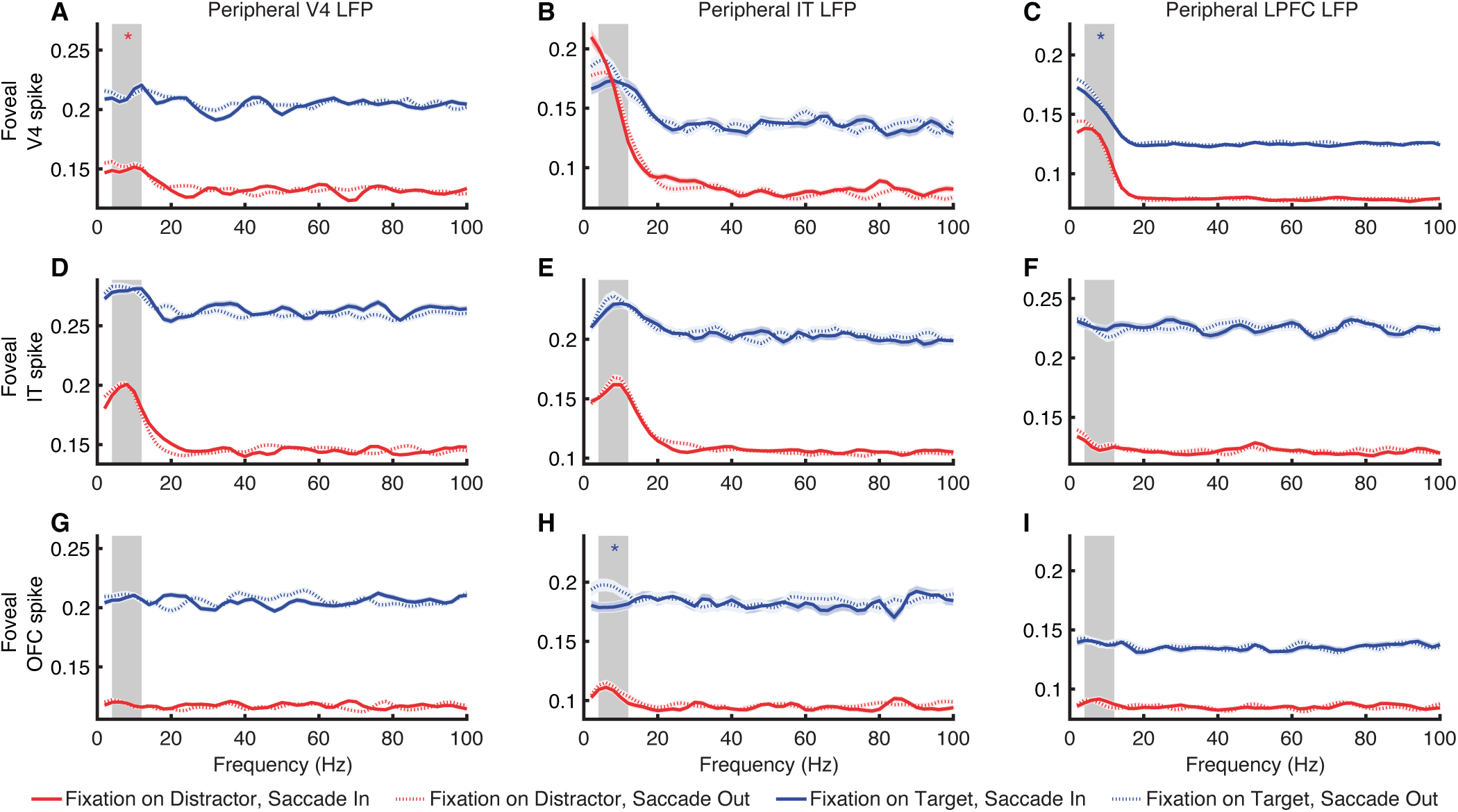
Spike-LFP coherence between foveal spikes (from units with a focal foveal receptive field) and peripheral LFPs for spatial attention. Legend conventions as in Fig. 6.

**Fig. S4.**
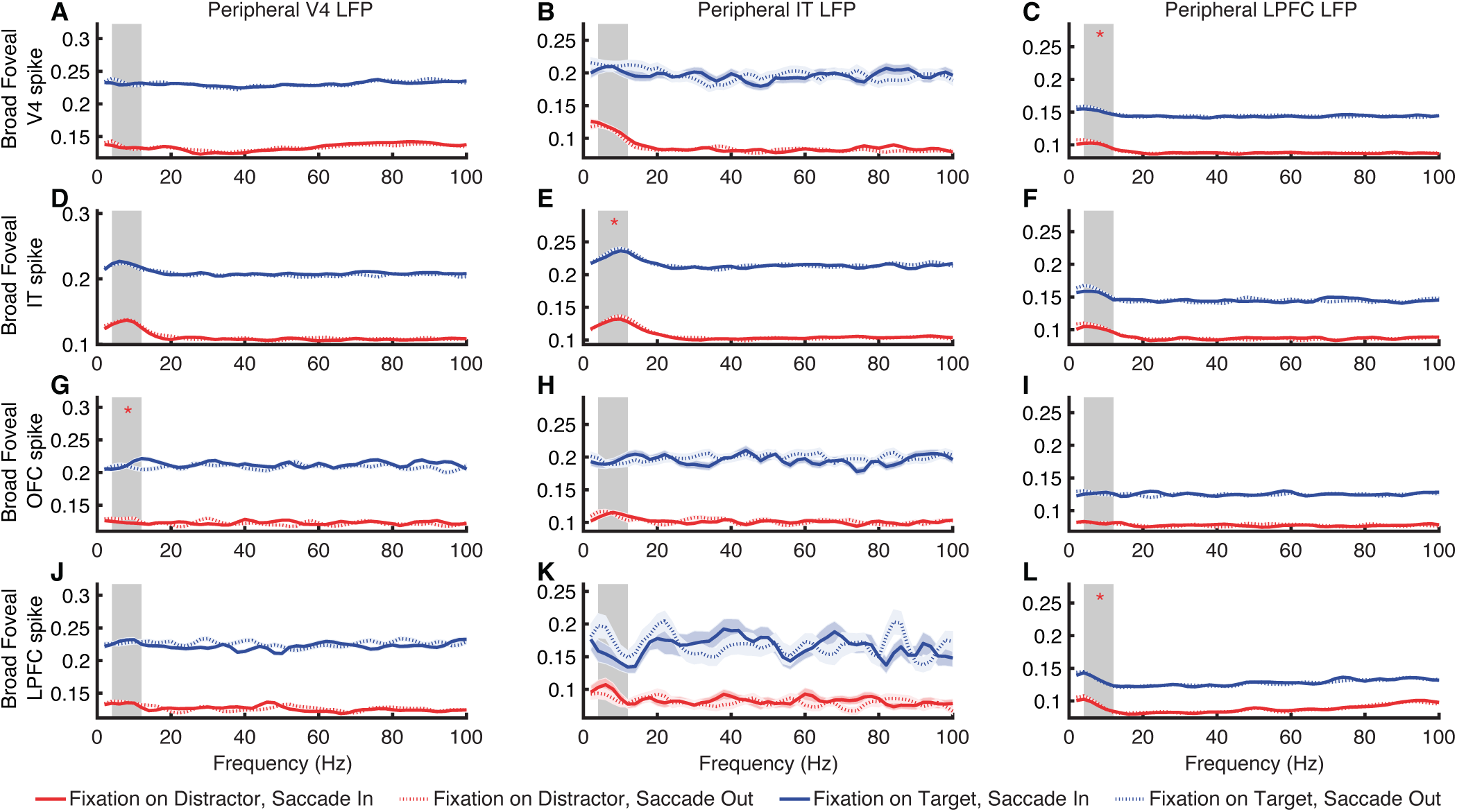
Spike-LFP coherence between foveal spikes (from units with a broad foveal receptive field) and peripheral LFPs for spatial attention. Legend conventions as in Fig. 6.

## References

1. Kastner, S. and L.G. Ungerleider, Mechanisms of visual attention in the human cortex, in Annual Review of Neuroscience. 2000. p. 315–341.

2. Corbetta, M. and G.L. Shulman, Control of goal-directed and stimulus-driven attention in the brain. Nat Rev Neurosci, 2002. 3(3): p. 201–15.

3. Petersen, S.E. and M.I. Posner, The Attention System of the Human Brain: 20 Years After. Annual Review of Neuroscience, 2012. 35(1): p. 73–89.

4. Moore, T. and M. Zirnsak, Neural Mechanisms of Selective Visual Attention. Annual Review of Psychology, 2017. 68(1): p. 47–72.

5. Fiebelkorn, I.C. and S. Kastner, Functional Specialization in the Attention Network. Annual Review of Psychology, 2020. 71(1): p. 221–249.

6. Wolfe, J.M., Guided Search 6.0: An updated model of visual search. Psychonomic Bulletin & Review, 2021. 28(4): p. 1060–1092.

7. Mazer, J.A. and J.L. Gallant, Goal-Related Activity in V4 during Free Viewing Visual Search: Evidence for a Ventral Stream Visual Salience Map. Neuron, 2003. 40(6): p. 1241–1250.

8. Bichot, N.P., A.F. Rossi, and R. Desimone, Parallel and Serial Neural Mechanisms for Visual Search in Macaque Area V4. Science, 2005. 308(5721): p. 529–534.

9. Burrows, B.E. and T. Moore, Influence and Limitations of Popout in the Selection of Salient Visual Stimuli by Area V4 Neurons. The Journal of Neuroscience, 2009. 29(48): p. 15169–15177.

10. Zhou, H. and R. Desimone, Feature-Based Attention in the Frontal Eye Field and Area V4 during Visual Search. Neuron, 2011. 70(6): p. 1205–1217.

11. Chelazzi, L., et al., A neural basis for visual search in inferior temporal cortex. Nature, 1993. 363(6427): p. 345–347.

12. Bichot, Narcisse P., et al., A Source for Feature-Based Attention in the Prefrontal Cortex. Neuron, 2015. 88(4): p. 832–844.

13. Zhou, H., Robert J. Schafer, and R. Desimone, Pulvinar-Cortex Interactions in Vision and Attention. Neuron, 2016. 89(1): p. 209–220.

14. Bichot, N.P., et al., The role of prefrontal cortex in the control of feature attention in area V4. Nature Communications, 2019. 10(1): p. 5727.

15. Buschman, Timothy J. and S. Kastner, From Behavior to Neural Dynamics: An Integrated Theory of Attention. Neuron, 2015. 88(1): p. 127–144.

16. Zhang, J., et al., Visual Attention in The Fovea and The Periphery during Visual Search. bioRxiv, 2021: p. 2021.11.22.469359.

17. Mendoza-Halliday, D., et al., Dissociable neuronal substrates of visual feature attention and working memory. Neuron, 2024. 112(5): p. 850–863.e6.

18. James, W., The principles of psychology, Vol I. The principles of psychology, Vol I. 1890, New York, NY, US: Henry Holt and Co. xii, 697-xii, 697.

19. Kinchla, R.A., Attention. Annual Review of Psychology, 1992. 43(Volume 43, 1992): p. 711–742.

20. Swallow, K.M. and Y.V. Jiang, Attentional Load and Attentional Boost: A Review of Data and Theory. Frontiers in Psychology, 2013. 4.

21. de Lissa, P., et al., In pursuit of visual attention: SSVEP frequency-tagging moving targets. PLOS ONE, 2020. 15(8): p. e0236967.

22. Yi, D.-J., et al., Neural fate of ignored stimuli: dissociable effects of perceptual and working memory load. Nature Neuroscience, 2004. 7(9): p. 992–996.

23. Lavie, N., Perceptual load as a necessary condition for selective attention. Journal of Experimental Psychology: Human Perception and Performance, 1995. 21(3): p. 451–468.

24. Lavie, N., Distracted and confused?: Selective attention under load. Trends in Cognitive Sciences, 2005. 9(2): p. 75–82.

25. Jyoti, M., et al., Neural Basis of Superior Performance of Action Videogame Players in an Attention-Demanding Task. The Journal of Neuroscience, 2011. 31(3): p. 992.

26. Morrone, M.C., V. Denti, and D. Spinelli, Color and Luminance Contrasts Attract Independent Attention. Current Biology, 2002. 12(13): p. 1134–1137.

27. Murphy, K.S.J. and J.A. Foley-Fisher, Visual search with non-foveal vision. Ophthalmic and Physiological Optics, 1988. 8(3): p. 345–348.

28. Bertera, J.H. and K. Rayner, Eye movements and the span of the effective stimulus in visual search. Perception & Psychophysics, 2000. 62(3): p. 576–585.

29. Cornelissen, F.W., K.J. Bruin, and A.C. Kooijman, The Influence of Artificial Scotomas on Eye Movements during Visual Search. Optometry and Vision Science, 2005. 82(1).

30. McIlreavy, L., J. Fiser, and P.J. Bex, Impact of Simulated Central Scotomas on Visual Search in Natural Scenes. Optometry and Vision Science, 2012. 89(9).

31. Nuthmann, A., On the visual span during object search in real-world scenes. Visual Cognition, 2013. 21(7): p. 803–837.

32. Treue, S. and J.C. Marínez-Trujillo, Feature-based attention influences motion processing gain in macaque visual cortex. Nature, 1999. 399(6736): p. 575–579.

33. Maunsell, J.H.R. and S. Treue, Feature-based attention in visual cortex. Trends in Neurosciences, 2006. 29(6): p. 317–322.

34. Fries, P., et al., Modulation of Oscillatory Neuronal Synchronization by Selective Visual Attention. Science, 2001. 291(5508): p. 1560–1563.

35. Jutras, M.J., P. Fries, and E.A. Buffalo, Oscillatory activity in the monkey hippocampus during visual exploration and memory formation. Proceedings of the National Academy of Sciences, 2013. 110(32): p. 13144–13149.

36. Yan, T. and H. Zhou, Synchronization between frontal eye field and area V4 during free-gaze visual search. Zoological Research, 2019. 40(5): p. 394.

37. Gross, C.G., How Inferior Temporal Cortex Became a Visual Area. Cerebral Cortex, 1994. 4(5): p. 455–469.

38. Logothetis, N.K. and D.L. Sheinberg, Visual object recognition. Annual Review of Neuroscience, 1996. 19(1): p. 577–621.

39. Tanaka, K., Mechanisms of visual object recognition: monkey and human studies. Current Opinion in Neurobiology, 1997. 7(4): p. 523–529.

40. Tsao, D.Y., et al., Patches of face-selective cortex in the macaque frontal lobe. Nature Neuroscience, 2008. 11: p. 877.

41. Rangel, A., C. Camerer, and P.R. Montague, A framework for studying the neurobiology of value-based decision making. Nat Rev Neurosci, 2008. 9(7): p. 545–556.

42. Rudebeck, P.H., et al., Effects of amygdala lesions on reward-value coding in orbital and medial prefrontal cortex. Neuron, 2013. 80(6): p. 1519–1531.

43. McGinty, Vincent B., A. Rangel, and William T. Newsome, Orbitofrontal Cortex Value Signals Depend on Fixation Location during Free Viewing. Neuron, 2016. 90(6): p. 1299–1311.

44. Xie, Y., C. Nie, and T. Yang, Covert shift of attention modulates the value encoding in the orbitofrontal cortex. eLife, 2018. 7: p. e31507.

45. Barbas, H., Organization of cortical afferent input to orbitofrontal areas in the rhesus monkey. Neuroscience, 1993. 56(4): p. 841–864.

46. Webster, M.J., J. Bachevalier, and L.G. Ungerleider, Connections of Inferior Temporal Areas TEO and TE with Parietal and Frontal Cortex in Macaque Monkeys. Cerebral Cortex, 1994. 4(5): p. 470–483.

47. Grimaldi, P., Kadharbatcha S. Saleem, and D. Tsao, Anatomical Connections of the Functionally Defined “Face Patches” in the Macaque Monkey. Neuron, 2016. 90(6): p. 1325–1342.

48. Xiao, W., et al., Feature-selective responses in macaque visual cortex follow eye movements during natural vision. Nature Neuroscience, 2024.

49. Tsao, D.Y., et al., A Cortical Region Consisting Entirely of Face-Selective Cells. Science, 2006. 311(5761): p. 670–674.

50. Yamins, D.L.K., et al., Performance-optimized hierarchical models predict neural responses in higher visual cortex. Proceedings of the National Academy of Sciences, 2014. 111(23): p. 8619.

51. Chang, L. and D.Y. Tsao, The Code for Facial Identity in the Primate Brain. Cell, 2017. 169(6): p. 1013–1028.e14.

52. Bashivan, P., K. Kar, and J.J. DiCarlo, Neural population control via deep image synthesis. Science, 2019. 364(6439): p. eaav9436.

53. Ponce, C.R., et al., Evolving Images for Visual Neurons Using a Deep Generative Network Reveals Coding Principles and Neuronal Preferences. Cell, 2019. 177(4): p. 999–1009.e10.

54. Bao, P., et al., A map of object space in primate inferotemporal cortex. Nature, 2020. 583(7814): p. 103–108.

55. Gross, C., et al., Inferior temporal cortex as a pattern recognition device, in Computational Learning and Cognition, E. Baum, Editor. 1993, Society for Industrial and Applied Mathematics: Philadelphia. p. 44–73.

56. Tanaka, K., Inferotemporal Cortex and Object Vision. Annual Review of Neuroscience, 1996. 19(1): p. 109–139.

57. Tsao, D.Y., et al., Faces and objects in macaque cerebral cortex. Nat Neurosci, 2003. 6(9): p. 989–995.

58. Wang, S., et al., Encoding of Target Detection during Visual Search by Single Neurons in the Human Brain. Current Biology, 2018. 28(13): p. 2058–2069.e4.

59. Wang, S., et al., Abstract goal representation in visual search by neurons in the human pre-supplementary motor area. Brain, 2019. 142(11): p. 3530–3549.

60. Bichot, N.P. and R. Desimone, Chapter 9 Finding a face in the crowd: parallel and serial neural mechanisms of visual selection, in Progress in Brain Research, S. Martinez-Conde, et al., Editors. 2006, Elsevier. p. 147–156.

61. Szczepanski, S.M., et al., Functional and structural architecture of the human dorsal frontoparietal attention network. Proceedings of the National Academy of Sciences, 2013. 110(39): p. 15806–15811.

62. Gregoriou, G.G., et al., High-Frequency, Long-Range Coupling Between Prefrontal and Visual Cortex During Attention. Science, 2009. 324(5931): p. 1207–1210.

63. Gregoriou, G.G., et al., Lesions of prefrontal cortex reduce attentional modulation of neuronal responses and synchrony in V4. Nature Neuroscience, 2014. 17(7): p. 1003–1011.

64. Stemmann, H. and W.A. Freiwald, Evidence for an attentional priority map in inferotemporal cortex. Proceedings of the National Academy of Sciences, 2019. 116(47): p. 23797–23805.

65. Thompson, K.G., et al., Perceptual and motor processing stages identified in the activity of macaque frontal eye field neurons during visual search. Journal of Neurophysiology, 1996. 76(6): p. 4040–4055.

66. Freiwald, W.A., D.Y. Tsao, and M.S. Livingstone, A face feature space in the macaque temporal lobe. Nat Neurosci, 2009. 12(9): p. 1187–1196.

67. Freiwald, W.A. and D.Y. Tsao, Functional Compartmentalization and Viewpoint Generalization Within the Macaque Face-Processing System. Science, 2010. 330(6005): p. 845.

68. Fries, P., et al., The Effects of Visual Stimulation and Selective Visual Attention on Rhythmic Neuronal Synchronization in Macaque Area V4. The Journal of Neuroscience, 2008. 28(18): p. 4823.

69. Saleem, K.S. and N.K. Logothetis, A combined MRI and histology atlas of the rhesus monkey brain in stereotaxic coordinates. 2012, London: Academic Press.

